# A high-throughput multiplexing and selection strategy to complete bacterial genomes

**DOI:** 10.1101/2021.06.14.448320

**Authors:** Sergio Arredondo-Alonso, Anna K. Pöntinen, François Cléon, Rebecca A. Gladstone, Anita C. Schürch, Pål J Johnsen, Ørjan Samuelsen, Jukka Corander

## Abstract

**Background:** Bacterial whole-genome sequencing based on short-read sequencing data often results in a draft assembly formed by contiguous sequences. The introduction of long-read sequencing technologies permits to unambiguously bridge those contiguous sequences into complete genomes. However, the elevated costs associated with long-read sequencing frequently limit the number of bacterial isolates that can be long-read sequenced.

Here we evaluated the recently released 96 barcoding kit from Oxford Nanopore Technologies (ONT) to generate complete genomes on a high-throughput basis. In addition, we propose a long-read isolate selection strategy that optimizes a representative selection of isolates from large-scale bacterial collections.

**Results:** Despite an uneven distribution of long-reads per barcode, near-complete chromosomal sequences (assembly contiguity = 0.89) were generated for 96 *Escherichia coli* isolates with associated short-read sequencing data. The assembly contiguity of the plasmid replicons was even higher (0.98) which indicated the suitability of the multiplexing strategy for studies focused on resolving plasmid sequences. We benchmarked hybrid and ONT-only assemblies and showed that the combination of ONT sequencing data with short-read sequencing data is still highly desirable: (i) to perform an unbiased selection of isolates for long-read sequencing, (ii) to achieve an optimal genome accuracy and completeness, and (iii) to include small plasmids underrepresented in the ONT library.

**Conclusions:** The proposed long-read isolate selection ensures completing bacterial genomes of isolates that span the genome diversity inherent in large collections of bacterial isolates. We show the potential of using this multiplexing approach to close bacterial genomes on a high-throughput basis.

## Introduction

Whole-genome sequencing (WGS) of bacterial isolates has dramatically increased its routine presence in clinical genomics and epidemiological investigations [1]. The possibility of using short-read technologies, affordable and generally accurate in terms of their sequencing reads, has permitted tracking the presence of particular sequencing types, assessing the presence of single-point mutations, or identifying antimicrobial resistance (AMR) genes in collections of thousands of bacterial isolates [2–4]. These applications are fundamental during outbreak detections and investigations, for which WGS has overcome some limitations associated with classical epidemiological techniques [2,5,6]. However, the read length associated with these short-read technologies (from 150 bp to 300 bp) cannot unambiguously span the presence of repeat elements such as insertion sequences (IS). This results in a fragmented assembly, typically formed by contiguous sequences (contigs) of unknown order. In particular, determining the genome context (chromosome, plasmid, phage) of genes typically surrounded by IS elements (e.g AMR genes) is challenging in draft assemblies [7,8].

Long-read sequencing technologies such as Oxford Nanopore Technologies (ONT) can generate genomic libraries with an average read length between 10-30 kbp [9], but with a lower associated raw read accuracy of ~97% (phred score ~15) depending on the pore chemistry [10]. These long-reads can typically span repeat elements in a bacterial genome producing a contiguous assembly consisting of single and circular contigs per replicon (chromosome and/or plasmids). However, the sequence accuracy of these complete genomes can suffer from incorrect basecalling of short homopolymer sequences resulting in early termination of open reading frames (ORF) in protein-coding genes [11].

An attractive alternative is to combine short- and long-read technologies to generate complete and accurate genomes in a process called hybrid assembly [12]. A frequent scenario encountered by researchers is that large collections of bacterial isolates have been massively short-read sequenced and only a subset of these isolates can be further selected for long-read sequencing. Therefore, selecting isolates for long-read sequencing is a non-trivial and crucial step that can affect subsequent analyses based on the resulting complete genomes.

There are different strategies to decrease the price of completing a genome with ONT sequencing [13,14]. Recently, Lipworth et al. [15] showed that usage of wash-kits coupled with shorter sequencing times can be optimized to obtain 36 complete genomes per single flow cell, achieving a reduction in long-read sequencing costs of 27%. During the hybrid assembly process, only a fraction of the total number of long-reads generated are required to bridge and span the initial short-read assembly graph and thus a low ONT coverage is sufficient to complete a genome.

The possibility of increasing the number of multiplexed isolates per flow cell is an alternative to the usage of wash-kits that reduces hands-on-time and avoids contamination issues between libraries. Recently, ONT has released a new barcoding kit allowing multiplexing 96 isolates in the same sequencing library.

Here, we evaluated the degree of completeness and accuracy in bacterial genomes generated using the new ONT barcoding kit for 96 bacterial isolates. In addition, we provide a step-by-step computational workflow to perform an unbiased selection of isolates for long-read sequencing based on the presence/absence of genes. as previously applied in two large bacterial collections of isolates [16,17].

The approach described in this study generated near-complete genomes for the majority of 96 *E. coli* isolates. The computational workflow proposed can be used to maximize and complete a representative selection of isolates from large-scale bacterial collections.

## Methods

### Long-read selection based on existing short-read assemblies

From a large collection of short-read isolates, frequently only a subset of isolates can be completed with long-read sequencing. We propose the following approach to select a representative subset of isolates for a population sample:

1. Define *M* as the presence/absence matrix of orthologous genes created by pangenome tools such as Roary [18] or Panaroo [19]. *M* is a binary matrix with *s* × *g* dimensions, in which *s* is the total number of isolates (samples) present in the collection and *g* corresponds to the total number of orthologous genes predicted.

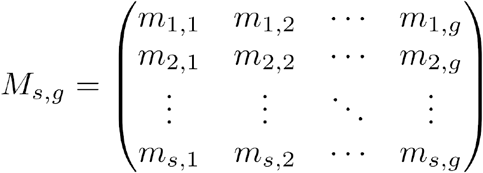
2. We transform *M* into a Jaccard distance matrix *D* with *s* × *s* dimensions using the R function *parDist* (method = ‘fjaccard’) provided in the R package parallelDist (version 0.2.4) https://github.com/alexeckert/parallelDist.

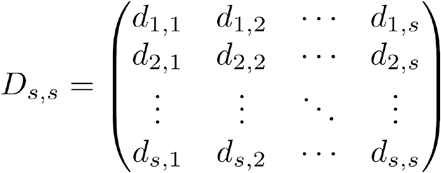 For instance, the element *d_1,2_* can be defined as the similarity distance between the genes predicted for the first isolate and second isolate:

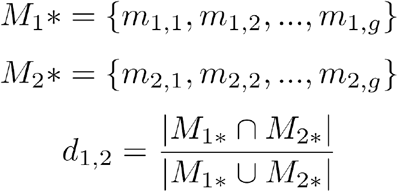
3. Next, the distance matrix *D* can be dimensionally reduced using the t-distributed stochastic neighbor embedding algorithm (t-sne) [20] which results in a new matrix *T* with only two dimensions. In this step, we use the R package Rtsne (version 0.15) https://github.com/jkrijthe/Rtsne considering as default a perplexity value of 30.
4. Next, we use the k-means algorithm [21] with *T* as an input data and define a number of centroids *c* such that it corresponds to the desired number of isolates to be long-read sequenced. A random initialization of the centroids permits defining the clusters *k* in *T* by assigning each point *t_i_* to its closest centroid. For each cluster *C_k_*, the algorithm updates the position of the centroid by computing the average Euclidean distance of each point *t_i_* against the centroid. The within-square variation *V* for a particular cluster *C_k_* can be defined as:

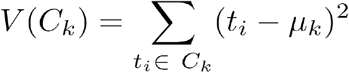 This process is repeated until convergence is reached, and thus the position of the centroids *c* in *T* no longer varies or the maximum number of iterations is reached (default = 1,000 iterations). To run the algorithm, we use the function *kmeans* provided in the R package stats (version 3.6.3). For a fixed number of centroids *c*, we run the k-means algorithm using 10,000 distinct initializations and select the initialization with the highest ratio between-cluster sum of squares and the total sum of squares (between_SS/tot_SS). This ratio is considered as the percent of total variance explained by the chosen number of centroids (clusters). The relationship between the percent of total variance explained and the number of centroids can be considered to visually determine the ideal numbers of clusters required to capture the diversity present in the collection (Elbow method).
5. For each cluster *C_k_*, we select for long-read sequencing the isolate *t_i_* with the lowest Euclidean distance with respect to its associated centroid *c*.

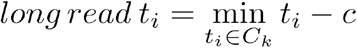
6. The tsne matrix *T* together with the final coordinates of the centroids *c* and the *t_i_* isolates selected for long-read sequencing are plotted using the R package ggplot2 (version 3.3.3).

The proposed workflow aims at capturing as comprehensive representation of the gene content variation across the lineages in the target population as possible and is available at as a https://gitlab.com/sirarredondo/long_read_selection Snakemake pipeline [22] that only requires (i) a presence/absence matrix in the same format as created by Roary/Panaroo and (ii) the desired number of long-read isolates.

### Collection of Illumina short-read assemblies

To showcase the proposed long-read selection, we considered the Norwegian *Escherichia coli* collection of 3254 isolates causing bloodstream infections recently described by Gladstone et al. [23]. This collection was short-read sequenced with the Illumina HiSeq platform and short-read contigs were created using VelvetOptimiser (version 2.2.5) https://bioinformatics.net.au/software.velvetoptimiser.shtml and Velvet (version 1.2.10) [24]. Furthermore, an assembly improvement step was applied to the assembly with the best N50 and contigs were scaffolded using SSPACE (version 2.0) [25] and sequence gaps filled using GapFiller (version 1.11) [26]. These short-read contigs will be later considered to compare the accuracy of the hybrid and ONT-only assemblies (section Genome accuracy and completeness).

The presence/absence of orthologous genes defined by Panaroo (version 1.0.2) [19] and PopPUNK lineages (version 2.0.2) [27] associated with the isolates were also extracted from Gladstone et al. [23] and considered as input for the long-read selection process. As an example, we fixed the number of centroids to 96 and considered the between_SS/tot_SS ratio to estimate if the maximum number of isolates that can be multiplexed in an ONT sequencer would suffice to capture the genomic diversity of this particular collection.

In the sections below, the number of centroids of the selection procedure were fixed to a large number of desired long-read isolates (n = 1085), of which we describe the ONT sequencing and analysis of 96 *E. coli* isolates.

### DNA isolation for ONT libraries

In total, 96 *E. coli* isolates, originally from clinical samples of human bloodstream infections, were grown on MacConkey agar No 3 (Oxoid Ltd., Thermo Fisher Scientific Inc., Waltham, MA, US) at 37°C, and individual colonies were picked for overnight growth in LB (Miller) broth (BD, Franklin Lakes, NJ, US) at 37°C in 700 rpm shaking. High-molecular-weight (HMW) genomic DNA from cell pellets of 1.6 ml overnight cultures was extracted using MagAttract^®^ HMW DNA Kit (Qiagen, Hilden, Germany) according to the manufacturer’s instructions and using a final elution volume of 100 μl. DNA concentration and integrity were verified using NanoDrop One spectrophotometer (Thermo Scientific) and the Qubit dsDNA HS assay kit (Thermo Fisher Scientific) on a CLARIOstar microplate reader (BMG Labtech, Ortenberg, Germany).

### ONT library preparation

The ONT library was prepared using SQK-NBD110-96 barcoding kit. Sequencing was run for 72 hours on GridION using FLO-MIN106 flow cells and MinKNOW v.20.10.6 software. Basecalling was conducted with the high-accuracy basecalling model and demultiplexing was performed by using Guppy v.4.2.3.

### Hybrid assemblies

Porechop (version 0.2.4) https://github.com/rrwick/Porechop was used to trim and remove ONT adapters with default parameters. Filtlong (version 0.2.0) https://github.com/rrwick/Filtlong was used to filter the ONT reads considering a minimum length of 1 kbp (--min_length 1000), a quality phred score of 20 (--mean_q_weight 20), retaining 90% of the total number of ONT reads (--keep_percent 90) from a maximum 40x coverage (--target_bases). Unicycler (version 0.4.7) [28] was run using the normal mode to perform a hybrid assembly with the Illumina trimmed reads and ONT reads retained after Filtlong.

We extracted the number of segments (contigs), links (edges), N50 and size of the components present at the resulting hybrid assembly graph. For each component, we defined its contiguity as:

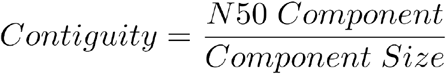

Mlplasmids (version 1.0.0) [29] was used with the *E. coli* model to confirm the origin (plasmid- or chromosome-derived) from the longest segment of each component.

### lllumina and ONT depth per replicon

To map the Illumina reads against each of the nucleotides assembled by Unicycler, we used Bowtie2 (version 2.4.2) [30] with the argument --very-sensitive-local, a minimum and maximum fragment length of 0 (-I 0) and 2000 respectively (-X 2000). The ONT reads were mapped against Unicycler assemblies using bwa mem (version 0.7.17) [31] and indicating the ready type (−x) as ont2d. Samtools (version 1.12) [32] was used with the commands sort, index and depth to process the alignments and retrieve the number of reads covering an individual nucleotide.

For each component present in the graph file given by Unicycler, we considered its longest contig and computed the average number of reads. To normalize the contig depth with respect to its chromosome, we divided the average depth of the contig against the average depth of the longest contig assembled by Unicycler (longest chromosome segment).

### ONT-only assemblies and long-read polishing

Flye (version 2.8.3-b1695) [33] was run with all the QC-passed ONT reads (phred score > 7) available per barcode (--nano-raw), specifying the option to recover plasmids (--plasmids), indicating an expected genome size of 5 Mbp (--genome-size 5m) and considering 3 polishing rounds (--iterations 3). For each component present in the assembly graph, we extracted the number of segments (contigs), links (edges), N50, component size and defined its component contiguity. Mlplasmids (version 1.0.0) [29] was used with the *E. coli* model to confirm the origin (plasmid- or chromosome-derived) from the longest segment of each component.

Following the recommendations to further polish Flye assemblies https://github.com/rrwick/Trycycler/wiki/Guide-to-bacterial-genome-assembly#13-medaka, we used Medaka (version 1.2.5) https://github.com/nanoporetech/medaka with the model ‘r941_min_high_g360’ and considered as input all the passed ONT reads available per barcode.

### Genome accuracy and completeness

Quast (version 5.0.2) [34] was used to compute alignments with a minimum size of 5 kbp (--min-alignment 5000) considering as reference short-read contigs assembled with Velvet (section Collection of Illumina short-read assemblies) from the same isolate against Unicycler, Flye and Medaka-polished contigs. For each reference contig, we kept the best alignment (--ambiguity-usage one). The average number of single-nucleotide polymorphisms (SNPs) and insertions and deletions (indels) per 100 kbp were considered as a reference-based metric of the genome accuracy shown by the hybrid and ONT-only assemblies.

For Unicycler assemblies, we also extracted the contigs present at the file ‘001_best_spades_graph.gfa’ corresponding to an optimal SPAdes assembly graph. Unicycler uses a range of k-mer sizes and computes a score between the resulting number of contigs and dead-ends to choose the SPAdes graph idoneal for the long-read bridge process. These SPAdes contigs were also compared using Quast against the reference contigs assembled by Velvet. This was relevant to determine the SNPs and indels per 100 kbp present at the SPAdes contigs. In this manner, we could assess if the accuracy (SNPs, indels) estimated for the hybrid assemblies was influenced by: (i) differences between the short-read assemblers, and/or (ii) regions of the genome from which its sequence is determined by the ONT reads, such as regions connecting dead-ends in the SPAdes graph.

The ideel test https://github.com/mw55309/ideel was used to obtain the number of early terminated ORFs in the assemblies [11]. Diamond (version 2.0.8) [35] was run with the blastp algorithm [36], specifying only a single target sequence per alignment (--max-target-seqs 1), a block size of 12 (−b12) and index chunk of 1 (−c1) against an index of the UniProt TREMBL database (retrieved in April 2021). For each hit, the ratio between query length and its own length was considered to assess the presence of interrupted ORFs (ratio < 0.9). This assesses whether the length of the predicted proteins is shorter than their closest hit, most likely caused by the introduction of a stop codon. Importantly, the total number of interrupted ORFs may include true pseudogenes, however, most of these hits are considered as non-true errors introduced due to a low sequencing accuracy. This ideel test provided a reference-free approach to assess the number of indels present in the final assemblies.

BUSCO (version 5.1.2) [37] with the genome mode (−m genome) and the lineage dataset (−l) ‘enterobacterales_odb10’ was used to assess the presence of 440 single-copy orthologous genes. BUSCO uses the following notation to annotate the genes as: complete (length within two standard deviations of the group mean length), fragmented (partially recovered) and missing (totally absent). The number of BUSCO complete genes was considered as a metric to assess the completeness of each assembly.

## Results

### A long-read selection spanning the genome diversity inherent in a short-read collection

In large bacterial collections for which possibly thousands of isolates are short-read sequenced, the selection of a subset of isolates for long-read sequencing is crucial to obtain a representative set of complete genomes spanning the genomic diversity present in the collection.

We propose the following approach summarised in the following steps (see Methods, for a formal description): i) consider the presence/absence matrix of orthologous genes computed by established pangenome tools such as Roary or Panaroo, ii) compute a distance matrix based on Jaccard distances, isolates are compared based on their shared number of orthologous genes, iii) reduce the dimensionality of the distance matrix using tsne by preserving local structure, iv) cluster the tsne dimensionally reduced matrix with the k-means algorithm, indicating as the number of centroids the desired number of long-read isolates and v) select the isolate with the closest Euclidean distance to its centroid. The proposed approach is fully available as a Snakemake pipeline at https://gitlab.com/sirarredondo/long_read_selection

We showcase the approach on a set of 3254 short-read sequenced isolates from a Norwegian longitudinal population genomic study of *E. coli* causing bacteraemia [23]. In this collection (Figure 1), we estimated that selecting 96 long-read isolates would capture ~ 99.9% of the total variance present in the tsne matrix computed from Jaccard distances.

**Figure 1.**
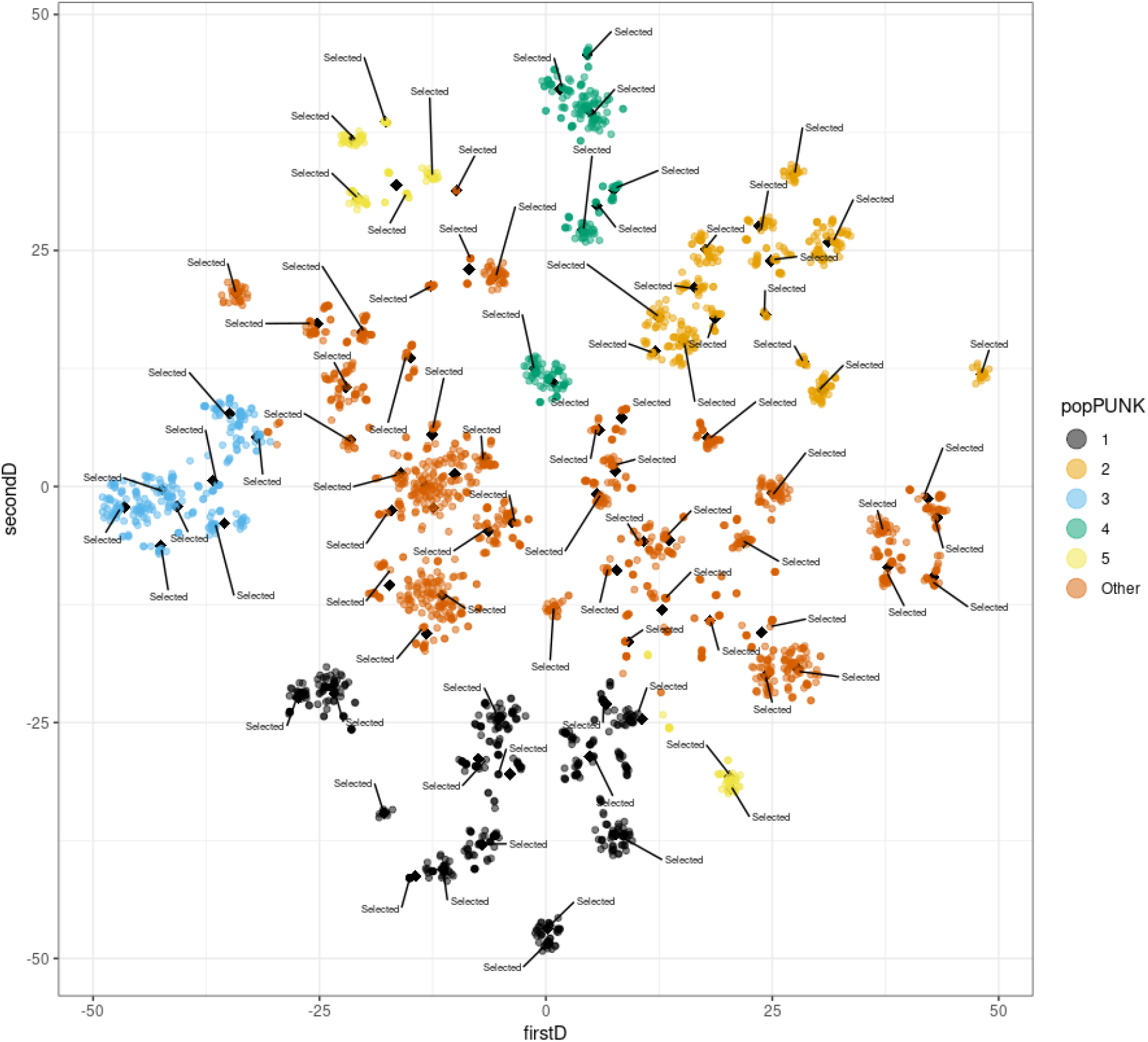
T-sne plot of the long-read selection, based on Jaccard distances computed from the presence/absence of genes defined by Panaroo in 3254 *E. coli* isolates. The five most predominant PopPUNK lineages are indicated with distinct colours, the rest of the lineages were merged into the category ‘Other’ (in yellow). The final coordinates of the centroids (n = 96) used in k-means are indicated as black diamond points. For each centroid (n = 96), the isolate closest to its respective centroid has been marked as ‘Selected’. In total, we indicate 96 *E. coli* isolates spanning the genomic diversity inherent in the collection that could be selected for further long-read sequencing.

### Uneven distribution of ONT reads in the 96 multiplexing approach

To investigate the possibility of obtaining complete genomes in a high-throughput manner, we evaluated the recently released ONT native barcode expansion kit to multiplex and sequence 96 *E. coli* isolates in the same MinION flowcell. These isolates belong to the Norwegian collection described above [23] for which short-read sequencing data are publicly available.

The ONT sequencing run generated a total of 10.71 Gbp QC-passed basecalled reads (average phred score > 7) that could be confidently assigned to a barcode, with an average N50 read length of 20.98 kbp (Figure 2). From the passed reads, the average ONT phred score corresponded to 12.48 equivalent to a read accuracy of 94.35 %. As previously reported for other multiplexing kits [13], we observed an uneven distribution of reads available per barcode which resulted in a large variation in the number of bases available per barcode (mean = 111.53 Mbp, median = 103.45 Mbp). This difference ranged from 3.37 Mbp (coverage - 0.67x) for barcode 68 to 244.00 Mbp (coverage - 48x) in barcode 73. Prior to the hybrid assemblies, ONT reads were filtered using Filtlong based on quality and length (see Methods), slightly reducing the number of reads available (mean = 94.36 Mbp, median = 85.39 Mbp). A complete description of ONT statistics per barcode is given in Suppl. Table S1.

**Figure 2.**
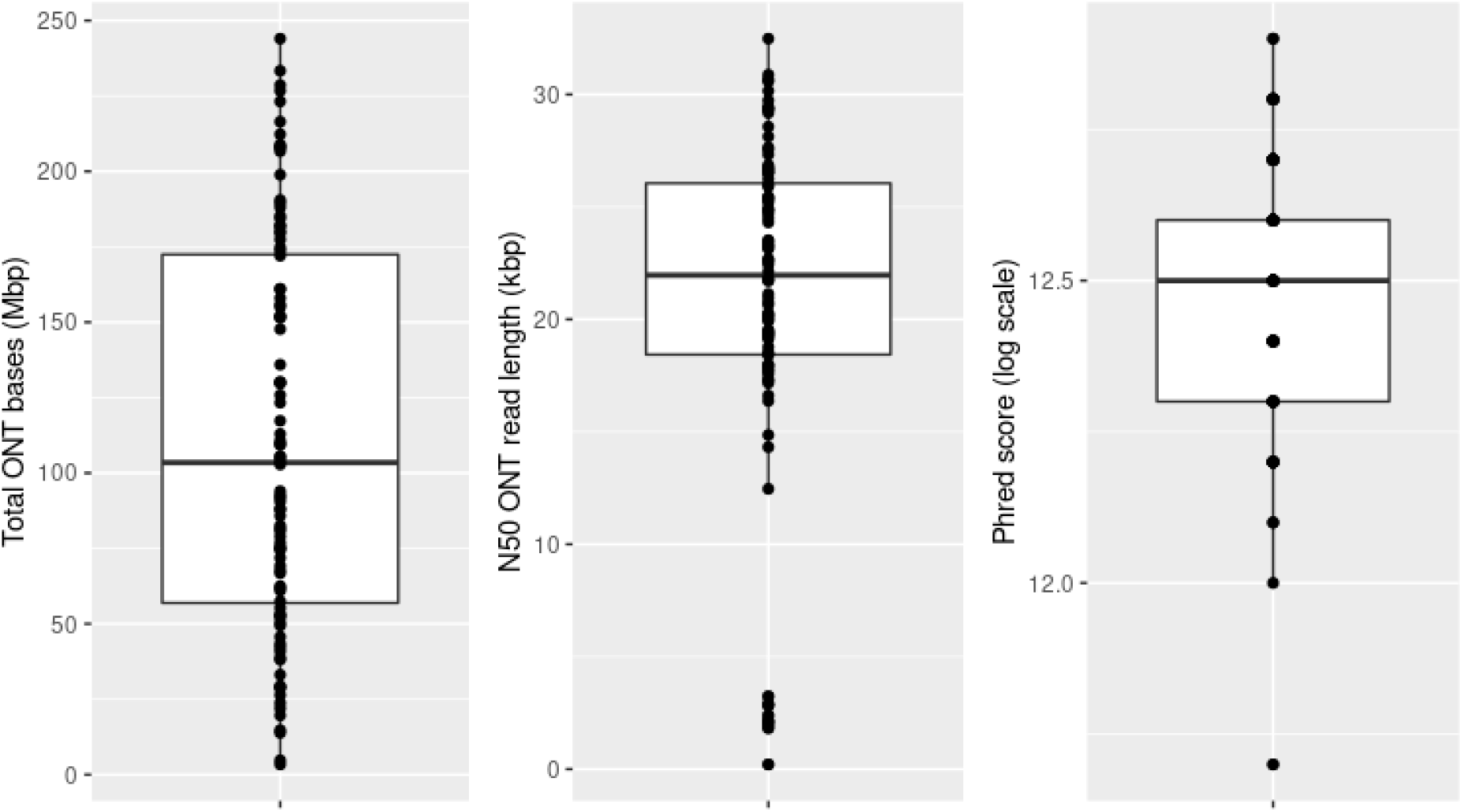
Oxford Nanopore Technologies (ONT) statistics per barcode (n = 96). First panel: boxplot of the number of bases (Mbp) generated in the sequencing run. Second panel: boxplot of the N50 ONT read length (kbp). Third panel: boxplot of the phred score (log10 scale) associated with the ONT reads.

### Evaluation of the hybrid assemblies

First, we focused on the chromosome sequence by analysing the largest component present in the hybrid assembly graph. In the 96 samples, the chromosome was present in a component with an average size of 5.0 Mbp (median = 5.03) and 11.39 contigs (median = 2.0) respectively (Figure 3). Despite the average number of contigs forming the chromosome, the contiguity (N50/component size) of the chromosome replicon was 0.89 (median = 1.0) which indicated that the chromosome component was assembled, for most samples, in a large contig (Figure 3). Furthermore, in 48 samples (50%) the chromosome resulted in a single circular contig (contiguity = 1.0) (Suppl. Table S2).

**Figure 3.**
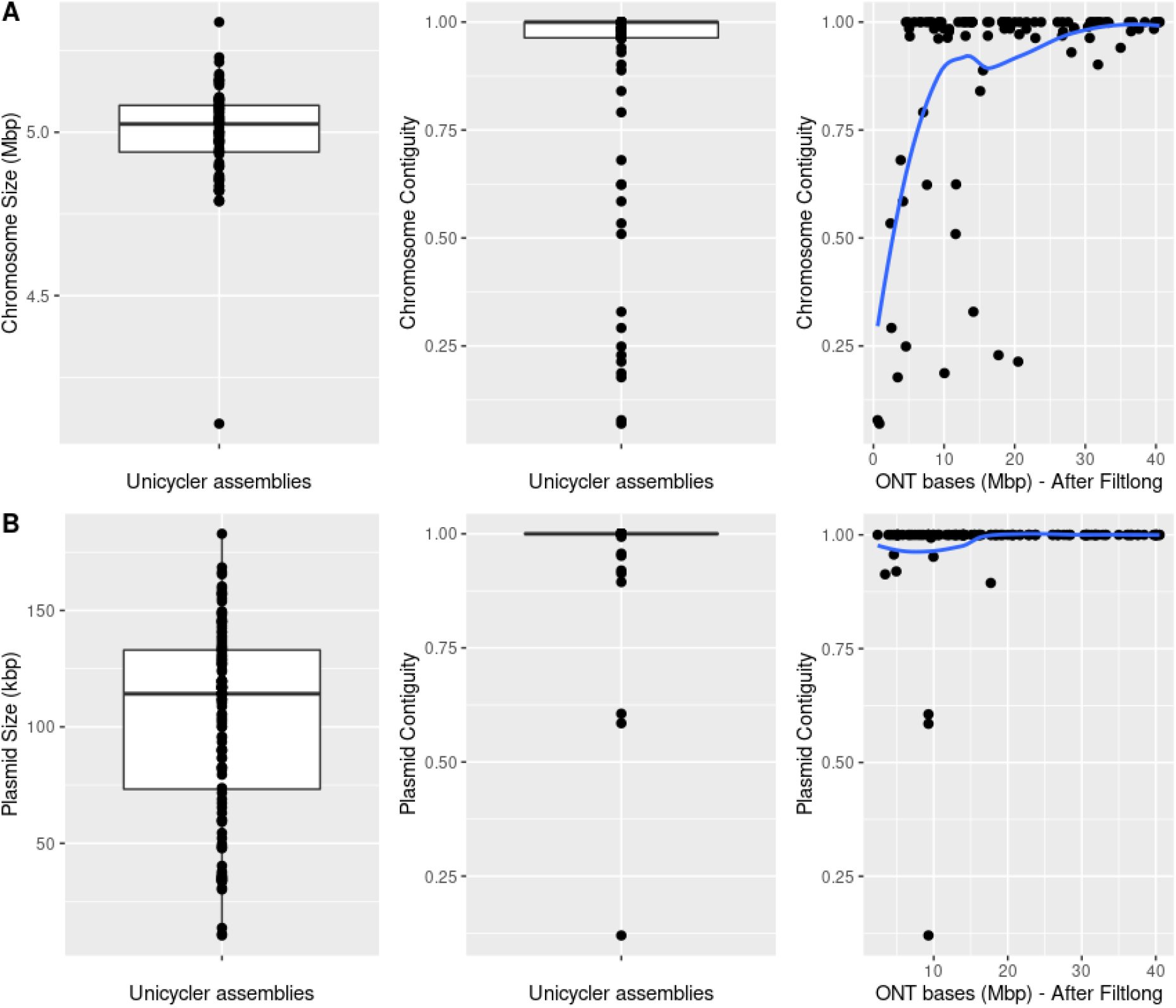
Unicycler statistics for the 96 isolates included in the ONT sequencing. Boxplots showing the component size (first panel) and contiguity values (second panel) together with an evaluation of the correlation between the number of ONT bases generated and the contiguity values achieved (third panel). A) Statistics for the chromosome component (largest component), the largest contig was confirmed as chromosome-derived with mlplasmids. B) Statistics for medium and large plasmid components (size > 10 kbp), the largest contig was confirmed as plasmid-derived with mlplasmids.

We observed a positive correlation (pearson corr. = 0.41) between the chromosome contiguity and the number of ONT reads available during the hybrid assembly (Figure 3). We assessed that an approximate ONT depth of ~5x was minimally required to achieve a perfect chromosome assembly (contiguity = 1.0). The lowest contiguity values corresponded to two isolates with a poor ONT coverage (< 1x) (barcode 65 and barcode 68). Overall, the isolates with an inferior contiguity value (threshold < 0.9) (18 barcodes, ~19%) had an associated low number of ONT reads available during the hybrid assembly (mean = 42.64 Mbp, coverage ~ 8.53x).

Next, we assessed the rest of the components present in the hybrid assembly graphs. The largest segment of each component was predicted with mlplasmids to confirm its plasmid origin. For medium and large plasmid components (size > 10 kbp) (n = 132), we obtained an average contiguity of 0.98 (median 1.0) and single circular contigs were retrieved for 118 components (89.4%) (Table 1). On average, the 132 plasmid components were formed by 1.79 contigs (median = 1.0). For the isolates with a low chromosome contiguity (< 0.9), the average plasmid contiguity was still 0.97 (median = 1.0) which indicated the suitability of the pipeline for plasmid reconstruction purposes. The correlation between plasmid contiguity and ONT reads available was weaker (pearson corr. = 0.16) than for the chromosomal component. Even for isolates with a poor ONT coverage, the contiguity values were close to 1.0 indicating that only a few long reads are sufficient to resolve plasmid components. The small plasmids (size < 10 kbp, n = 78) had an average contiguity of 0.99 (median = 1.0) and single circular contigs were retrieved for 74 plasmids (94.9%). The absence of repeat sequences in small plasmids permits to already obtain circular replicons by only using short-read sequencing assemblies and thus ONT reads are not required.

### Underrepresentation of small and medium plasmids in the ONT library

For each component present in the Unicycler assemblies, we determined its Illumina-read and ONT-read relative depth considering the average chromosomal depth as basis for the normalization, as previously performed [13]. This analysis was fundamental to confirm whether small plasmids were underrepresented in the ONT library preparation, as recently reported for ONT ligation kits [38].

As shown in Figure 4, small plasmids (size < 10 kbp) were strongly underrepresented in the ONT read output (log2 −0.63) in contrast to the Illumina read output (log2 4.79). For 7 small plasmids, we did not observe any ONT reads covering these replicons. These plasmids are usually present in high copy numbers in the cell as a survival and inheritance mechanism [39]. Therefore, we expected that these replicons would be overrepresented in the sequencing output resembling the plasmid read output given by Illumina (Figure 4). For medium plasmids (from 10 to 50 kbp), we also observed the same trend even though the underrepresentation in the ONT library was less pronounced (log2 −0.40). Notably, we observed the opposite trend for large plasmids (> 50 kbp) for which there was an underrepresentation in the Illumina library (log2 −0.60) compared to the ONT library (log2 0.18).

**Figure 4.**
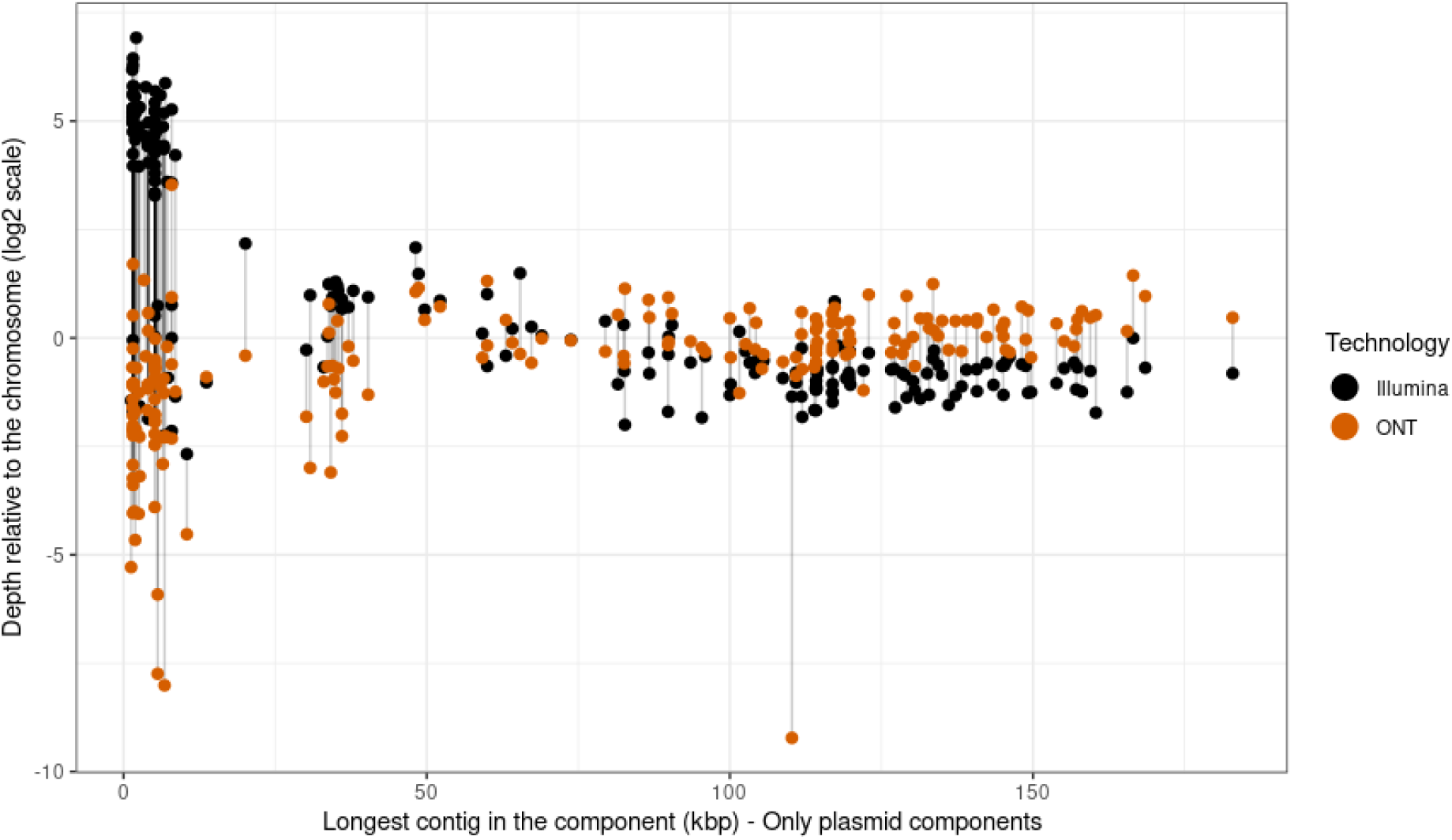
Illumina and ONT depth relative coverage of the plasmid components present in the Unicycler assemblies. The average depth of the plasmid replicons was normalized against the average depth of the chromosome to obtain a relative depth (log2 scale). Each plasmid is represented by two connected points depending on the sequencing technology.

### ONT-only assemblies

To evaluate whether Illumina reads are still required to obtain accurate genomes, we used Flye and Medaka to perform and polish assemblies based only on ONT reads. The largest component in the assembly graph had an average size of 4.45 Mbp (median = 4.98 Mbp) and 1.16 contigs (median = 1.0) (Suppl. Table S3). Flye failed to produce an assembly for 2 isolates (barcode 65 and barcode 68) for which the number of ONT reads generated was below 5 Mbp (Suppl. Table S1). In 71 samples (76%), the chromosome was represented by a single circular contig. The average chromosomal contiguity was 0.98 (median = 1.0) which indicated a higher contiguity with respect to the hybrid assemblies. However, the average size of the largest component (mean = 4.45 Mbp, median = 4.98 Mbp) was shorter than for Unicycler assemblies (mean = 5.0 Mbp, median = 5.03 Mbp). This indicated that for some isolates the size of the largest component did not match with the expected replicon size (~ 5Mbp).

The analysis of the rest of the components present in Flye assemblies (n = 346) revealed a high number of replicons with a chromosome origin (n = 193) indicating fragmentation of the chromosome into several components. For the components with a plasmid origin (n = 153), we obtained single circular contigs for 120 plasmids (78.43%). We observed a clear difference between the hybrid and ONT-only assemblies with respect to small plasmids (size < 10 kbp). Only 12 small circular plasmids were recovered in the Flye assemblies, in comparison to the 78 small plasmids present in the hybrid assembly. The absence of these plasmids in the Flye assemblies could be explained by their underrepresentation or complete absence in the total ONT read output, as shown in the section above.

### Genome accuracy and completeness of hybrid and ONT-only assemblies

To compare the accuracy of the resulting genomes, the hybrid assemblies and Flye assemblies were compared in terms of SNPs and indels considering both reference-based and reference-free methodologies (see Methods). Despite the fact that Flye incorporates a consensus-error module to correct the resulting genome sequences, we polished the Flye assemblies using Medaka and compared the genome accuracy against hybrid (Unicycler) and stand-alone Flye assemblies.

The hybrid assemblies showed the best accuracy stats with an average of 6.93 SNPs/100 kbp (median = 5.24) and 0.31 indels/100 kbp (median = 0.21), considering as ground truth non-repetitive alignments (> 5 kbp) against short-read contigs generated with Velvet. The SPAdes assemblies created by Unicycler with only short-reads showed an average of 0.85 SNPs/100 kbp (median = 0.28) and 0.06 indels/100 kbp (median = 0.04). This indicated that the accuracy of the hybrid assemblies was affected by the incorporation of error-prone ONT reads into the final genome sequence. The existence of dead-ends in the initial SPAdes graph indicated that parts of the genome were not sequenced with short-reads, thus their sequence had to be completed with long-reads and could not be polished with Illumina reads. As shown in Figure S2, two barcodes (69 and 72) had an elevated number of dead-ends (38 and 43) which resulted in a high number of SNPs/100 kbp (barcode 69: 23.89; barcode 72: 33.36) and indels/100 kbp (barcode 69: 1.25; barcode 72: 1.32).

Flye assemblies exhibited a higher number of errors with an average of 130.17 SNPs/100 kbp (median = 14.68) and 140.82 indels/100 kbp (median = 41.52). We observed a negative correlation between the number of ONT bases generated and the resulting number of SNPs (pearson corr. = −0.27) and indels (pearson corr. = −0.61) (Figures 5A and 5B). The correction performed by Medaka on Flye assemblies showed a reduction in the number of indels (mean = 126.35, median = 22.84) but the number of mismatches remained similar (mean = 130.29, median = 12.57). Again, we observed a negative correlation between the ONT depth and the resulting number of SNPs (pearson corr. = −0.27) and indels (pearson corr. = −0.61) (Figures 5A and 5B). For instance, for barcode 73 with the highest ONT read depth, the ONT-only assemblies had a number of SNPs (Flye = 6.52 SNPs/100 kbp, Medaka = 6.25 SNPs/100 kbp) comparable to Unicycler results (5.58 SNPs/100 kbp) which indicated that an increase in the ONT read depth can be highly beneficial to correct SNP errors. However, in the case of indels, we still observed 26.47 indels/100 kbp and 6.46 indels/100kbp for Flye and Medaka assemblies respectively, which is far from Unicycler results (0.04 indels/100 kbp).

**Figure 5.**
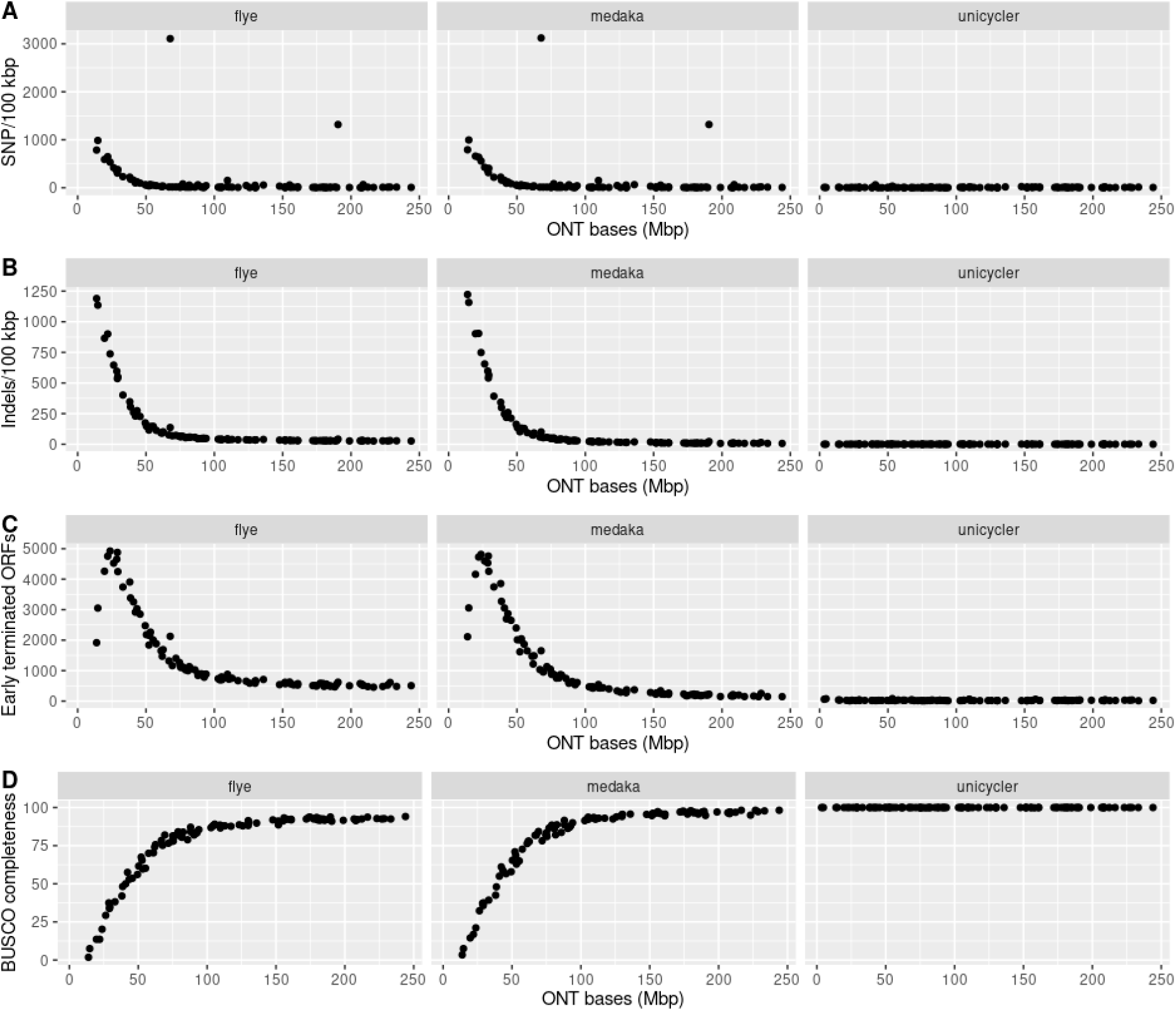
Correlation between the number of ONT bases generated per isolate and genome accuracy statistics for Flye assemblies (first panel), Medaka-polished assemblies (second panel) and Unicycler assemblies (third panel). A) Number of indels per 100 kbp computed considering as reference short-read contigs generated by an independent assembler. B) Number of SNPs per 100 kbp computed considering as reference short-read contigs generated by an independent assembler. C) Number of early terminated ORFs based on the ideel test. D) Completeness of the assemblies as percentage of complete single-copy conserved BUSCO genes (n = 440).

Following on this, we considered a reference-free approach to evaluate the impact of non-corrected indels in the interruption of ORFs using the ideel test [11]. In the hybrid assemblies, the number of promptly terminated ORFs corresponded to 25.07 (median = 21.0) which may include true pseudogenes. For Flye assemblies, on average the number of interrupted ORFs was 1380.64 (median = 772.5) and the correction with Medaka improved the genome accuracy by reducing the number to 1127.99 interrupted ORFs (median = 470.0). We observed a negative correlation between the number of ONT bases available and the number of interrupted ORFs present in Flye assemblies (pearson corr. = −0.76) and Medaka polished assemblies (pearson corr. = −0.78) (Figure 4C). We observed a plateau, around ~500 and ~150 ORFs for Flye and Medaka polished assemblies, for which an increase in ONT depth did not translate into a lower number of early terminated ORFs. Even for the isolate with the highest ONT depth (barcode 73, ONT bases 244 Mbp), the number of interrupted ORFs was still 146 for the Medaka polished assembly in comparison to 23 early terminated ORFs for the hybrid assembly.

Lastly, we evaluated the completeness of the genomes searching for complete orthologous genes against a curated set of 440 *Enterobacterales* single-copy conserved genes (BUSCO genes). For the hybrid assemblies, we recovered all BUSCO genes (100%) in contrast to only 76.6% for Flye assemblies. Furthermore, the percentage of missing genes was 10.3% which indicated that the *E. coli* genomes assembled by Flye were not complete. Medaka correction increased the number of recovered BUSCO genes (mean = 80.5%) and slightly decreased the number of missing genes (mean = 9.3%). For Flye and Medaka assemblies, a strong positive correlation of 0.78 was observed for both strategies between the ONT depth and the number of BUSCO complete genes (Figure 4D). The completeness of the ONT assemblies also stabilized despite an increase in the ONT read depth, around ~92% and ~98% for Flye and Medaka assemblies.

## Discussion

In this study, we have shown that near-complete genomes can be retrieved by multiplexing 96 bacterial isolates in the same long-read sequencing run. Despite an uneven distribution of ONT reads per barcode, we observed an average chromosomal contiguity of 0.89 which indicated that for most samples the chromosome was mostly represented by a single long contig. Furthermore, 92% of the plasmid sequences were circularized even for isolates with a low ONT read depth which makes this high-throughput multiplexing strategy an attractive choice for plasmid studies.

The availability of short-read sequencing data from the same bacterial isolates is still strongly desirable for three main reasons: (i) to perform an unbiased selection of isolates for long-read sequencing, (ii) to rely on the sequence accuracy achieved by short-read technologies (phred score > 30) and (iii) to include small plasmids underrepresented in the ONT library.

We proposed that given a short-read collection of isolates, the pangenome of these isolates can be computed and the presence/absence matrix of orthologous genes considered as the basis for long-read selection. This selection ensures that complete genomes are generated from distinct clusters which are defined by the gene content of the isolates present in the collection. Furthermore, the generated complete genomes can be used to conduct reference-based approaches relying on the short-read data of the non-long read sequenced isolates belonging to a particular genomic cluster. A limitation of this approach is that the selected isolate may not possess other accessory genes present in clonally related isolates. This is particularly relevant for medium/small plasmids or phage elements consisting of only a few genes and thus having a small relative weight in the distance matrix used as a basis to assign the clusters.

We observed that an increase in the ONT read depth is critical to improve the accuracy of ONT-only assemblies, in particular for SNP calling. However, in the case of indels, the systematic and non-random ONT read errors reported in homopolymer sequences resulted in a high number of early terminated ORFs, even for the isolates with the highest ONT read depth (~ 150 ORFs). Given the uneven distribution of ONT reads observed in the multiplexing approach, the accuracy and completeness of ONT-only assemblies for the isolates with a low read depth can result in a high number of SNPs and indels in their complete genome sequences. This drawback still allows to identify the genomic context from a gene-of-interest (e.g AMR genes) but can limit studies based on SNP signatures such as outbreak investigations.

In the hybrid assemblies, the accuracy of the complete genomes is unaffected by the ONT read depth since, in general, the long-reads are only used as bridges to unequivocally connect short-read contigs. With fewer long-reads, we could still obtain a complete genome and its accuracy was determined by the error read associated with the short-read technology. However, the accuracy of the hybrid assemblies can be affected by the quality of the initial short-read graph. If there is an elevated number of dead-ends in the short-read assembly graph, the sequence of ONT reads would be considered to complete the resulting genome and polishing that particular genomic region with Illumina reads would not be possible.

As previously reported, we also observed an underrepresentation of small plasmids in the ONT library which resulted in their absence in ONT-only assemblies. These plasmids can carry AMR or virulence genes and overlooking their presence in ONT-only assemblies when using ligation kits can affect subsequent analyses [38]. Small plasmids, however, are present in the initial short-read graph, usually as single circular contigs, and thus the absence of ONT reads covering these replicons do not affect the true representation of the genome.

In conclusion, we have shown the potential of using the recently released ONT nanopore barcoding kit for 96 bacterial isolates to recover near-complete genomes in combination with prior short-read sequencing data. We propose a long-read isolate selection based on the gene content to ensure that the resulting complete genomes span the diversity present in the collection. Finally, the possibility of generating complete genomes on a high-throughput basis will likely continue to significantly advance the field of microbial genomics.

## Data availability

ONT and Illumina sequencing data are available through the ENA Bioprojects PRJEB45354 and PRJEB32059 respectively. For each ONT barcode, individual ONT and Illumina accessions are indicated in Table S1. ONT reads and assemblies are available through the following permanent Figshare datasets: ONT reads -https://doi.org/10.6084/m9.figshare.14778333; Unicycler assemblies - https://doi.org/10.6084/m9.figshare.14705979; Flye assemblies - https://doi.org/10.6084/m9.figshare.14706015; Medaka polished assemblies - https://doi.org/10.6084/m9.figshare.14706108. The long-read selection pipeline is available at https://gitlab.com/sirarredondo/long_read_selection. An Rmarkdown document with the code and files required to reproduce the results presented in this manuscript is available at https://gitlab.com/sirarredondo/highthroughput_strategy

## Additional Files

Table S1. Summary of the ONT read statistics and ENA read accessions.

Table S2. Assembly statistics and mlplasmids prediction of the components present in the Unicycler assembly graph.

Table S3. Assembly statistics and mlplasmids prediction of the components present in the Flye assembly graph.

## Abbreviations

AMR: Antimicrobial Resistance
INDEL: Insertions and deletions
IS: Insertion Sequence
ONT: Oxford Nanopore Technologies
ORF: Open-Reading-Frame
SNP: Single-Nucleotide Polymorphism
WGS: Whole-Genome Sequencing

## Author’s contributions

PJJ, ØS and JC designed and sought funding for the study. SA-A, AS and JC designed and implemented the long-read selection pipeline. AKP and FC performed the HMW DNA extractions. SA-A performed the computational analyses: read processing, assembly and genome statistics. RAG facilitated the short-read sequencing data and assemblies, and provided the popPUNK lineages. SA-A, AKP and JC wrote the first draft of the manuscript. All authors contributed and reviewed the manuscript.

## Acknowledgments

We thank Lars Håland and Christopher Fenton at the Genomics Support Center Tromsø for generating, basecalling and demultiplexing the ONT sequencing reads.

## Supplementary Figures

**Figure S1.**
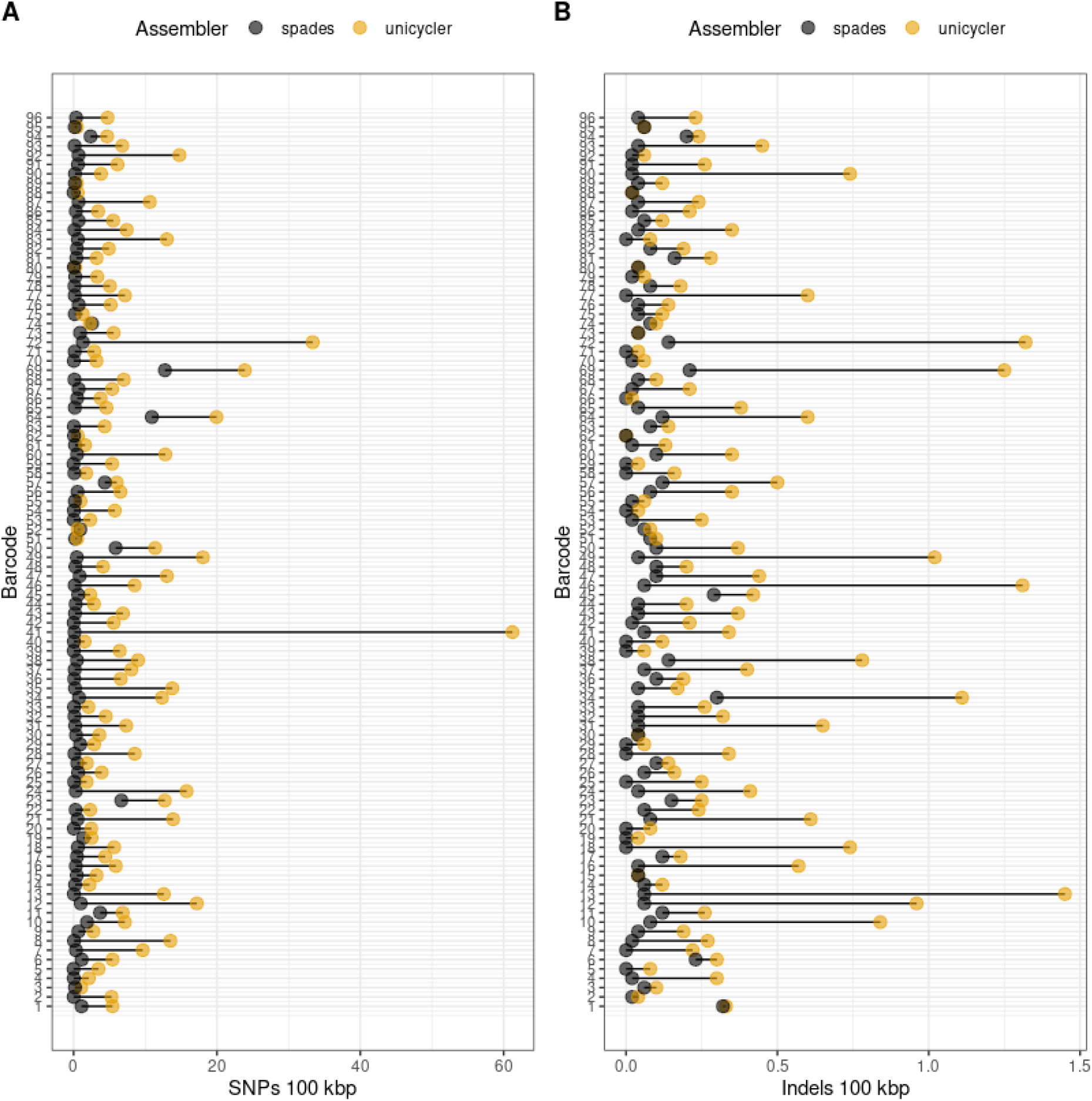
Genome accuracy comparison between SPAdes assemblies and Unicycler (hybrid) assemblies considering as reference short-read contigs assembled with Velvet. A) Number of SNP differences per 100 kbp observed in the SPAdes assemblies (black circles) and Unicycler assemblies (yellow circles). For each barcode, the difference in SNPs/100 kbp is represented by a line connecting the results of the two assemblies. B) Number of indel differences per 100 kbp observed in the SPAdes assemblies (black circles) and Unicycler assemblies (yellow circles). For each barcode, the difference in indels/100 kbp is represented by a line connecting the results of the two assemblies.

**Figure S2.**
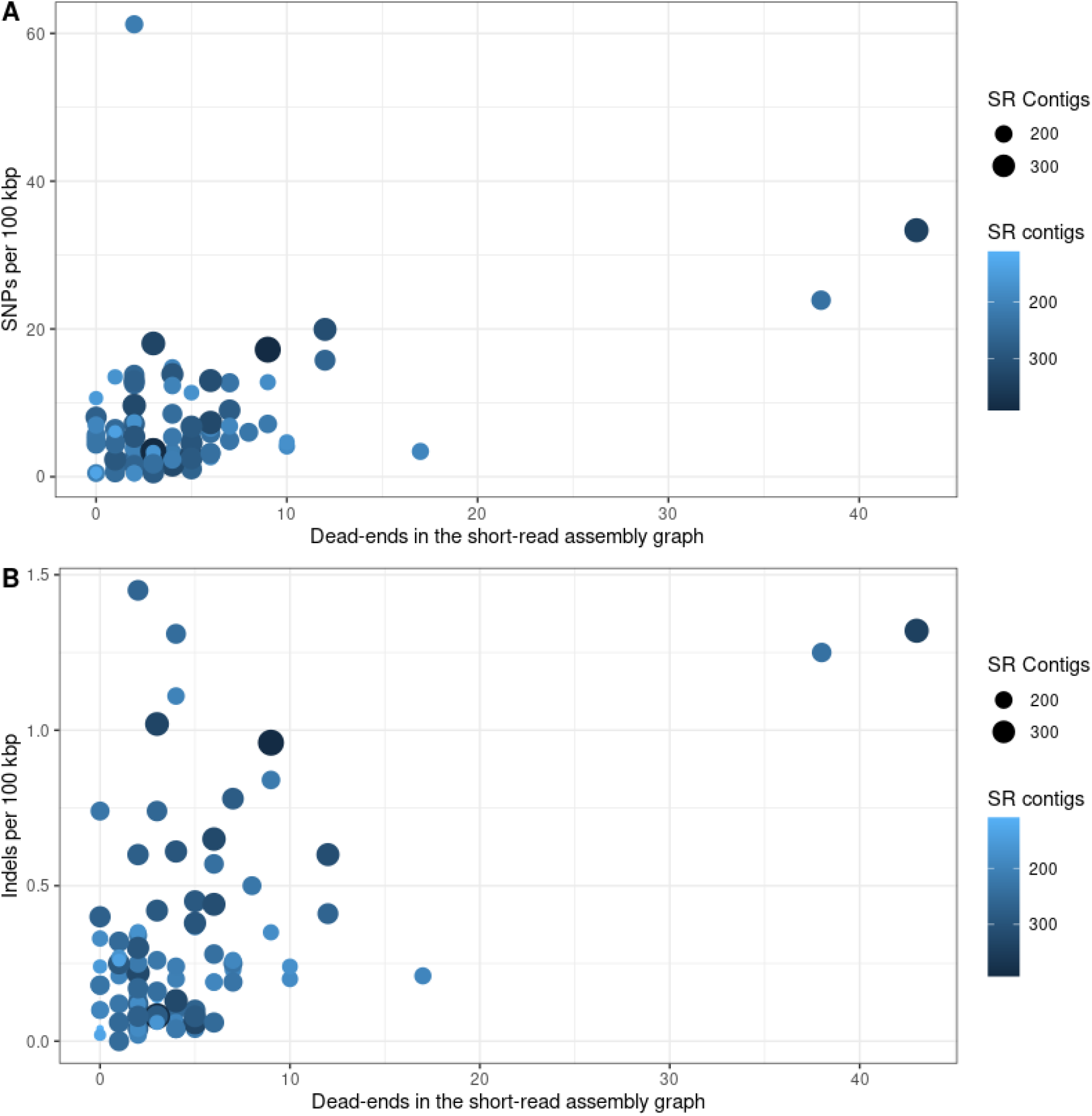
Overview of the hybrid assembly accuracy based on the quality of the initial SPAdes assembly graph in terms of dead-ends (x-axis). A) Number of SNP differences per 100 kbp in comparison to the number of dead-ends present in the SPAdes assembly graph. Each dot represents a barcode and its size and colour differs depending on the number of contigs (SR contigs) present at the SPAdes assembly graph. B) Number of indel differences per 100 kbp in comparison to the number of dead-ends present in the SPAdes assembly graph. Each dot represents a barcode and its size and colour differs depending on the number of contigs (SR contigs) present at the SPAdes assembly graph.

